# Forest degradation reshapes trophic functioning in vertebrate food webs across Amazonian forests

**DOI:** 10.64898/2026.07.13.737465

**Authors:** Ana Carolina Antunes, Ulrich Brose, Anelise Montanarin, Benjamin Rosenbaum, Carlos A. Peres, Carsten Meyer, Henrique Miguel Pereira, Jessica Hines, Jingyi Li, Tainara Sobroza, Emilio Berti, Adrian Barnett, Alexine Keuroghlian, Antonio Carlos da Silva Zanzini, Arlison B. Castro, Benoit de Thoisy, Carlos Rodrigo Brocardo, Clarissa Rosa, Daniel da Silva Ferraz, Daniel Gomes da Rocha, Dian Carlos Pinheiro Rosa, Diogo Maia Gräbin, Eduardo Nakano-Oliveira, Elildo Alves Ribeiro de Carvalho, Eloisa Neves Mendonça, Emerson M. Vieira, Emiliana Isasi-Catalá, Emiliano Esterci Ramalho, Fabricio Baccaro, Fernanda Michalski, Fernanda Santos, Fernando Anaguano Yancha, Francesca Belem Lopes Palmeira, Gabriel de Avila Batista, Galo Zapata-Rios, Gilson de Souza Ferreira Neto, Guilherme Costa Alvarenga, Helena Alves do Prado, Hugo C M Costa, João Marcelo Deliberador Miranda, Julia Salvador Maldonado, Karen Borges-Almeida, Leonardo Maracahipes-Santos, Lucas Paolucci, Luciana Zago da Silva, Maíra Benchimol, Marcela Guimarães Moreira Lima, Paula Ribeiro Prist, Paulo Monteiro Brando, Raphael Foscarini, Renato Richard Hilário, Ricardo Sampaio, Robert B. Wallace, Rossano Marchetti Ramos, Rodolfo Vasquez Martínez, Samir G. Rolim, Rocio del Pilar Rojas Gonzáles, Santiago Espinosa, Tony Enrique Noriega Piña, Wagner Tadeu Vieira Santiago, Ana Yoko Ykeuti Meiga, Ana Paula de Almeida Correa, André Pinassi Antunes, Victor Lery Caetano-Andrade, Regison da Costa de Oliveira, Benoit Gauzens

**Author notes:** Correspondence and requests for materials should be addressed to Ana Carolina Antunes.

## Abstract

The Amazon is a mosaic of ecosystems, accounting for 13% of all known species globally, and responsible for providing a variety of ecosystem services, including a key role in global climate regulation. However, 18% of the Amazon forest cover has been lost completely and a further 38% degraded, threatening biodiversity and millions of local livelihoods. Currently, little is known about how this impacts the functioning of ecological communities. Here, we combine an energetic approach with biodiversity metrics to quantify ecosystem functioning in mammal and bird food webs across Amazonia, along a gradient of forest degradation linked to road proximity. We show that even under relatively low disturbance, ecosystem functions shift: carnivory increases closer to roads, driven by generalist species that persist under these conditions, whereas herbivory declines mainly due to reduced herbivore biomass and species richness. This suggests that processes associated with forest degradation can alter energy flow even where biodiversity metrics remain relatively unchanged, highlighting energetic approaches as sensitive indicators of ecosystem disruption.

## 1. INTRODUCTION

Tropical forests regulate climate, sustain vast biodiversity, and provide essential ecosystem services to human societies ^1^. The persistence of these services depends on the maintenance of ecosystem functioning, which is increasingly threatened by human-driven environmental changes ^2^. Across Amazonian forests, such alterations are driven primarily by two distinct yet interacting processes: deforestation and forest degradation ^3^. Deforestation involves the conversion of forest into non-forest land cover and, in Brazil, is concentrated along the so-called “arc of deforestation” ^4^. In contrast, forest degradation involves a progressive decline in forest integrity through edge effects, fire, selective logging, and droughts ^3^. Although often less visible than deforestation, degradation now affects a comparable, or even larger, extent of Amazonian forests, with far-reaching consequences for biodiversity and ecosystem processes ^3,5^.

Forest degradation is strongly driven by the expansion of road networks across tropical forests. This network increases human access to previously intact forests, accelerating settlement, resource extraction, land conversion and introduction of non-native species and diseases ^6^. In turn, roads promote the proliferation of formal and informal access networks that penetrate forest interiors, creating the characteristic “fishbone” deforestation pattern. Both paved and unpaved roads are associated with forest degradation, by increasing access to even remote protected areas, triggering a range of direct and indirect impacts such as roadkills ^7^, expansion of selective logging ^8^, increased edge effects and fragmentation, declines in habitat quality and increased fire susceptibility ^4,9^. Even within Indigenous territories, the expansion of formal and informal road networks has been linked with increasing deforestation rates ^10^. Substantial ecological degradation can occur even where deforestation remains limited, as road expansion facilitates resource extraction, hunting and other diffuse patterns of disturbance that are difficult to detect and monitor at large spatial scales ^11,12^. Deforestation also increases human access to wildlife, intensifying overhunting and defaunation ^13^, both recognized as major forms of forest degradation ^12,14^. Consequently, the cumulative spread of both formal and informal secondary roads has increased landscape accessibility across the Amazon, acting as a proxy for multiple anthropogenic pressures, including forest degradation, habitat fragmentation, hunting, and land-use change, with persistent consequences for biodiversity and ecosystem functioning ^15,16^.

Beyond altering habitat structure, roads also have direct and indirect effects on vertebrate communities that play key roles in ecosystem functioning and ecosystem services delivery ^17^. These impacts can occur even in areas with relatively low road density, showing that road effects extend far beyond their immediate physical footprint ^18^. However, the consequences for species richness and community composition vary across taxa, often favoring smaller, generalist and omnivore species ^11,17,19^ while having strong negative impacts on apex predators and larger-bodied forms ^20,21^. Ultimately, changes in community structure can reshape ecosystem functioning by altering predator–prey dynamics and energy flow across trophic levels ^22^ (Fig. 1).

**Fig. 1:**
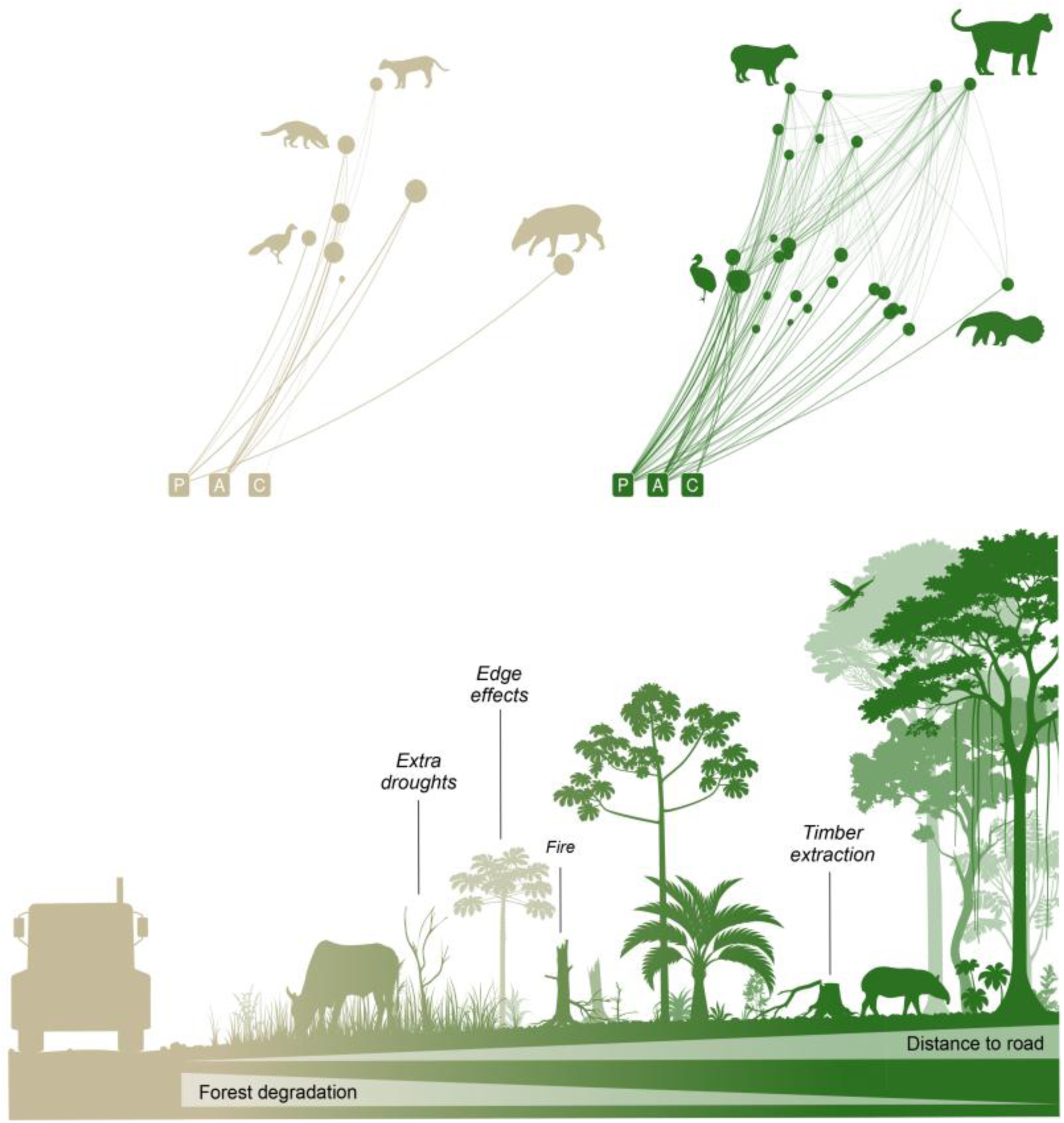
Road-associated forest degradation reshapes energy fluxes across Amazonian ecological networks. The lower panel illustrates a conceptual gradient of increasing forest degradation associated with proximity to roads, including ecological disturbances such as edge effects, drought intensification, fire, and timber extraction. Upper panels represent local ecological networks distributed along this gradient, and associated energy fluxes. Basal resource compartments are represented by P = plants, A = other animals (including invertebrates and small vertebrates not detected by camera traps), and C = carrion. Nodes represent species, and node sizes are proportional to species biomass, while link thickness and transparency reflect the relative magnitude of energy fluxes between trophic levels. Species silhouettes are illustrative and represent only a subset of the community.

Despite growing evidence that forest degradation and deforestation alter species composition and abundance, their consequences for ecosystem functioning remain poorly understood, particularly at broad spatial scales across tropical forests. A mechanistic approach to address this gap is provided by the bioenergetic framework, which integrates food-web structure, metabolic theory, and biodiversity data to quantify ecosystem functions such as herbivory and carnivory through energy fluxes across ecological networks ^23,24^. Here, we hypothesised that road proximity would primarily reduce carnivory and herbivory fluxes indirectly, by reshaping community biomass and reducing species richness and body-size structure, particularly through the loss of large-bodied species, thereby propagating changes in energy flux through food webs.

We used the largest camera-trap dataset assembled across Amazon forests ^25^, comprising 3,613 camera traps distributed across 86 spatial clusters (Fig. 2) that recorded 107 species over 19 years, enabling a macroecological approach to translate biodiversity data into ecosystem functions. We quantified energy fluxes as proxies of carnivory and herbivory by homeotherm vertebrates (mammals and birds) to assess how these key ecosystem functions respond to proximity to roads across the Amazon. Carnivory was defined as the consumption of animal prey, including both vertebrates and invertebrates, as smaller carnivores feed extensively on invertebrates, while herbivory was the consumption of plant-based resources ^26,27^. Because road networks are major drivers of both deforestation and forest degradation, we used distance to the nearest road as an integrative proxy for human disturbance across the study sites. By linking biodiversity structure to trophic energy flow, we provide a basin-wide assessment of how forest degradation induced by road proximity reshapes vertebrate trophic functioning. Because energy flux represents a common currency linking organisms and ecosystem functions across trophic levels ^24^, it provides a robust pathway to model how changes in diversity will affect ecosystem functions at large spatial scales, while capturing underlying ecological processes ^24,28,29^. This is particularly relevant for understanding resilience and recovery potential in vast tropical forest regions, which is essential to inform conservation and restoration strategies.

**Fig. 2:**
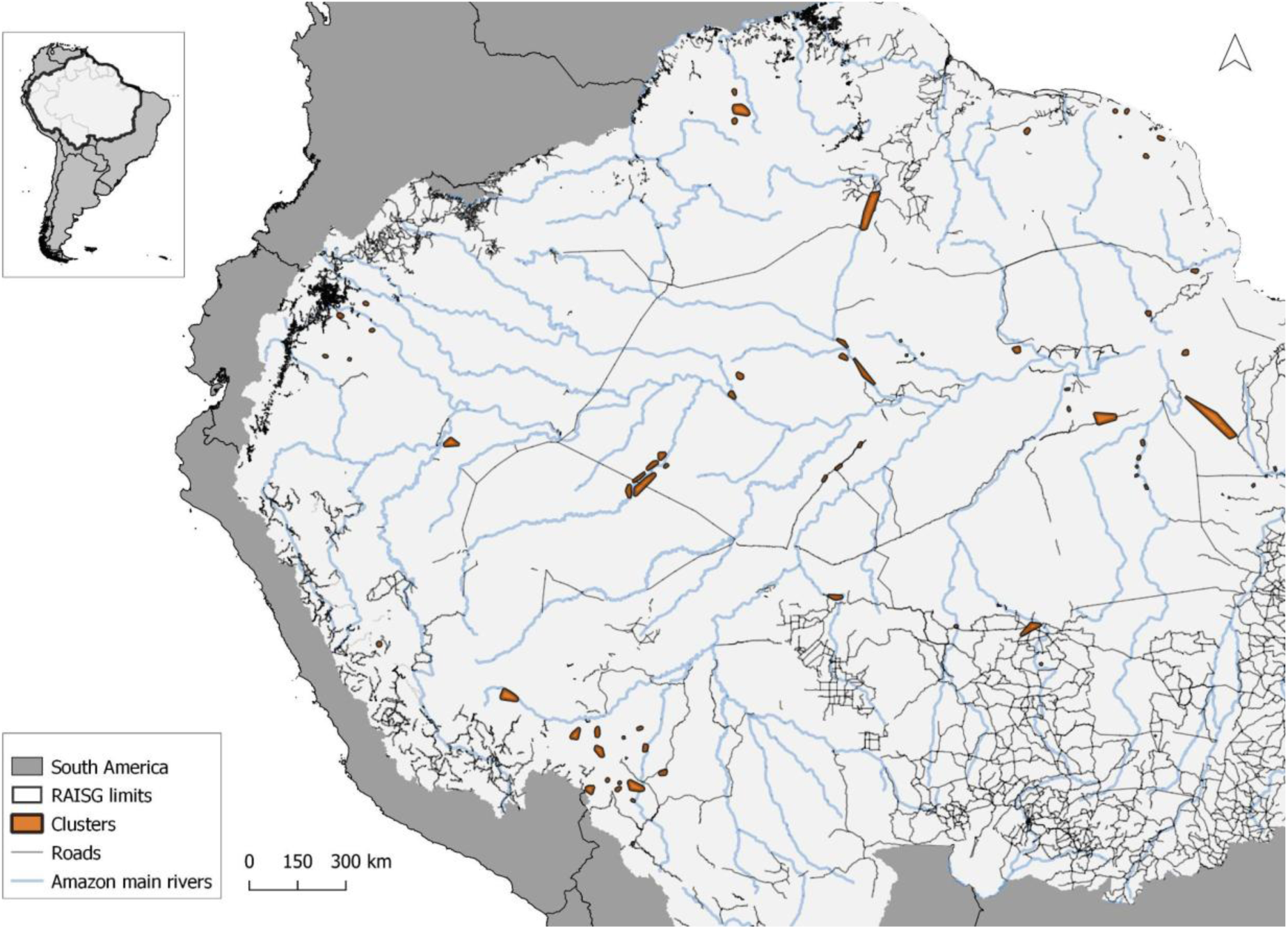
Spatial distribution of study sites across Amazonia. Map of camera-trap clusters (orange) sampled across the Amazon within the phytogeographic and hydrological boundaries of Amazonia (RAISG). Roads are shown in grey and major rivers in blue.

## 3. RESULTS

After excluding one outlier, the final analysis included 85 camera-trap clusters (i.e. local food webs), across which we detected 35 bird and 72 mammal species. Based on dietary composition, 78 and 84 species contributed to carnivory and herbivory fluxes, respectively, whereas 56 species contributed to both trophic pathways. To assess how road proximity influences carnivory and herbivory across the Amazon, both directly and indirectly through its effects on community structure, we fitted two Structural Equation Models (SEMs).

For carnivory, the final SEM (Fig. 3a) showed an adequate model fit (Fisher’s C = 7.85, p = 0.13). Carnivory fluxes increased with carnivore biomass (standardized coefficient β = 0.29, p = 0.014) and carnivore species richness (β = 0.55, p < 0.001), while mean body mass had no detectable effect (β = 0.05, p = 0.50). Importantly, distance to roads had a direct negative effect on carnivory flux (β = −0.21, p = 0.020) (Fig. 4), indicating higher levels of carnivory nearer roads after accounting for richness, biomass, and body mass. Carnivore biomass increased strongly with species richness (β = 0.79, p < 0.001) and decreased with mean body mass (β = −0.15, p = 0.023), while road distance had no detectable effect on either carnivore species richness (β = 0.29, p = 0.10) or mean body mass (β = −0.15, p = 0.24).

**Fig. 3:**
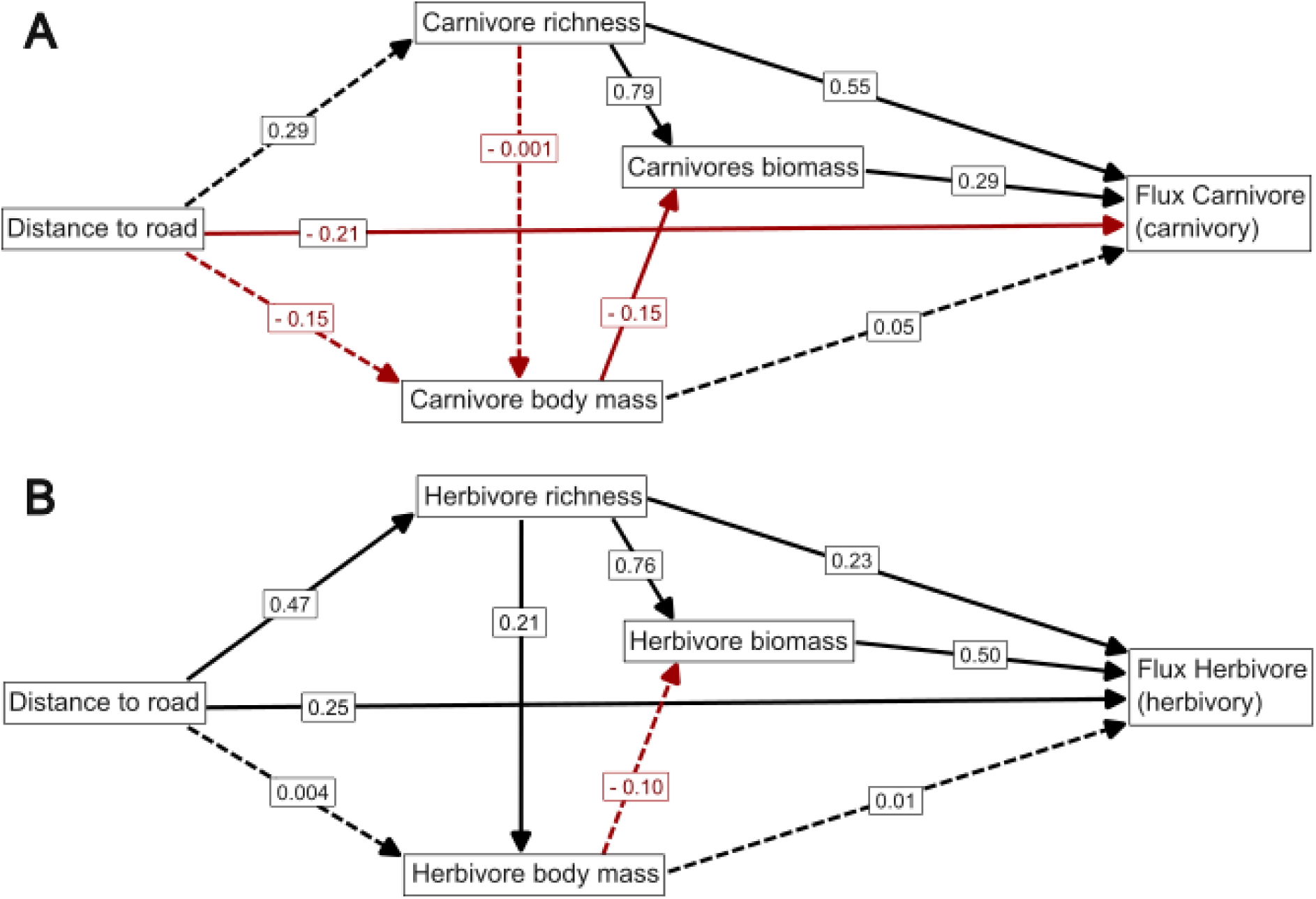
Structural equation model linking road proximity, trophic community properties and energy flux (ecosystem functions). Path diagrams showing relationships between distance to roads, species richness, mean body mass, total biomass and energy flux for (a) carnivores (carnivory flux) and (b) herbivores (herbivory flux). Solid lines indicate significant paths and dashed lines non-significant paths with their standardized path coefficients. Red arrows highlight negative effects, whereas black arrows represent positive relationships.

**Fig 4.**
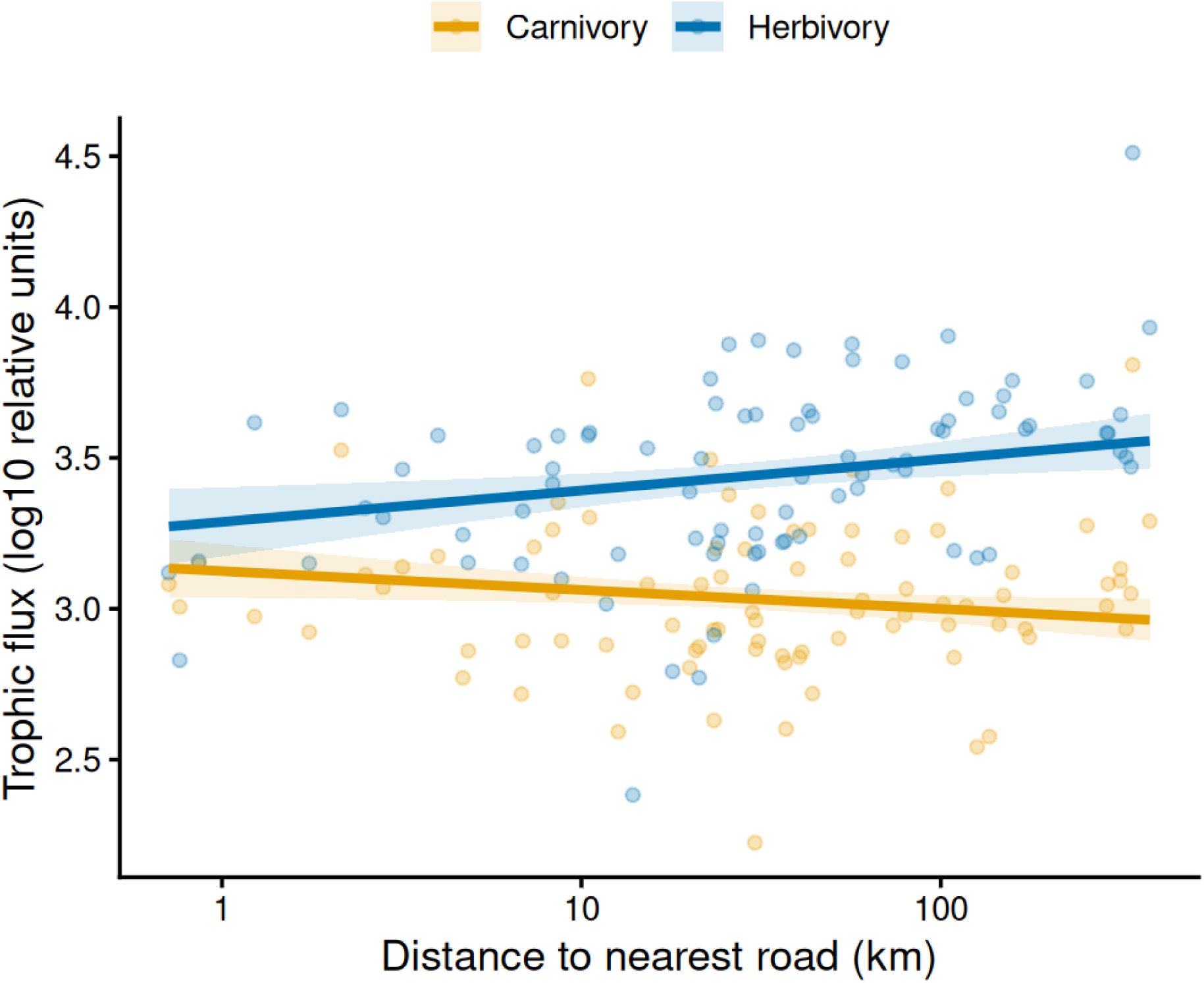
Direct effects of road distance on carnivory and herbivory fluxes. Data points show observed log₁₀-transformed trophic flux (relative units) for each camera-trap cluster (N= 85) as a function of distance to roads (on a log_10_ scale). Lines represent SEM predictions while holding community biomass, mean body mass, and species richness at their mean values. Shaded areas show 95% confidence intervals around the predicted effects.

For herbivory, the final SEM (Fig. 3b) showed a good model fit (Fisher’s C = 5.53, p = 0.25). Herbivory fluxes increased strongly with herbivore biomass (β = 0.50, p < 0.001) and, to a lesser extent, herbivore species richness (β = 0.23, p = 0.046), whereas mean body mass had no detectable effect (β = 0.01, p = 0.89). Distance to roads showed a positive direct association with herbivory flux (β = 0.25, p = 0.008) (Fig. 4), indicating higher herbivory fluxes in areas farther from roads. Road distance also increased herbivore species richness (β = 0.47, p = 0.010), which strongly increased herbivore biomass (β = 0.76, p < 0.001), supporting an indirect positive effect of road distance on herbivory fluxes through community structure. Herbivore mean body mass increased slightly with species richness (β = 0.21, p = 0.044) but was unrelated to road distance (β = 0.004, p = 0.97). Mean body mass showed no detectable relationship with herbivore biomass (β = −0.10, p = 0.17).

Results were robust to the exclusion of a potential outlier cluster in Yasuní National Park, Ecuador, located near a low-traffic road associated with oil and gas operations. Further analyses with and without this cluster yielded consistent results, and we therefore present the full dataset in the Supplementary Information (Section 1). Together, these models reveal opposite responses to road proximity, with carnivory intensified near roads, whereas herbivory increased with distance from roads, while energy fluxes in both trophic pathways were primarily structured by community biomass and species richness.

## 4. DISCUSSION

In this study, we integrated tropical forest biodiversity surveys using camera traps with metabolic and network theory to quantify modelled energy fluxes (i.e. relative estimates of trophic energy transfer) and link them to key ecosystem functions mediated by terrestrial birds and mammals, including herbivory and carnivory, at a Pan-Amazonian scale. By combining biodiversity data with energetic estimates, our approach provides the first large-scale, spatially explicit assessment of how human disturbances reshape vertebrate trophic processes in Amazonian forests. Even across landscapes experiencing relatively low to moderate human pressure, including many of the sampled sites, we detected measurable shifts in inferred trophic functioning, particularly herbivory and carnivory fluxes mediated by camera-trap-detectable medium- to large-bodied birds and mammals. Specifically, carnivory increased closer to roads, whereas herbivory showed the opposite pattern, mediated by reductions in herbivore richness and biomass. These patterns indicate that distance to roads reshape trophic processes by unevenly affecting species abundances and biomass distributions according to their sensitivity to disturbance. More broadly, our findings show that the energetic structure of ecological communities can reveal early functional disruption, even where some conventional biodiversity metrics remain stable, exposing changes in ecosystem functioning that would otherwise remain undetectable through species richness, abundance, or biomass estimates alone.

Areas closer to roads are typically associated with edge effects, including canopy opening, shifts in plant composition and altered resource availability ^30,31^. Although vertebrate responses to these changes remain poorly understood across the Amazon, local studies suggest that species differ strongly in their tolerance to disturbance: generalists often persist or even increase under low-intensity habitat modification, whereas specialists and species with narrow diet-niche breadth tend to decline ^32,33^. In our low-impact study areas, however, disturbance levels appear insufficient to reduce carnivores, which is consistent with our finding that predator species richness, biomass, and mean body size did not vary with road proximity. Nonetheless, modelled carnivory flux increased near roads, indicating that predator communities are likely functionally reorganized even when their taxonomic structure remains unchanged. This pattern is likely driven by few generalist predators (Supplementary Fig 1), such as tayras, opossums, coatis, and some small felids, whose mobile, opportunistic, and often omnivorous, traits enable them to exploit elevated resource availability near forest edges, including increases in fruits, insects, and small vertebrates resulting from canopy gaps and microclimatic changes ^34,35^. In addition, large apex predators may remain less affected under lower levels of habitat degradation due to their often-wide-ranging movements, which allow them to track resources across broader landscapes even when local habitat conditions and/or prey base deteriorate ^21^. Together, these patterns suggest that stable levels of species richness and biomass can mask important functional turnover within assemblages ^36^, where a smaller subset of tolerant generalists may contribute disproportionately to trophic functioning as more sensitive specialists decline, amplifying carnivory fluxes near roads.

For herbivores, the mechanisms underlying road effects appear to differ markedly from those shaping carnivores. Previous studies have shown that road networks in the Amazon can significantly alter herbivore communities, affecting both species’ richness and community composition, with declines occurring not only in disturbance-sensitive specialists but also in several generalist taxa ^11^. Human-driven changes in ecological communities can alter the flow of energy from plants to vertebrates ^37^, with our results showing reduced herbivory flux in proximity to roads driven by both direct and indirect drivers. Road-adjacent habitats may support herbivore assemblages that are functionally reorganized, with more disturbance-sensitive or specialized species declining while more generalist species persist (Supplementary Fig. 1). However, these structural changes did not fully explain the observed pattern, suggesting that road proximity may alter trophic functioning beyond changes in species richness, biomass, or body size alone. Roads can disrupt interspecific interactions and reorganize trophic pathways ^38^, potentially reducing the diversity and relative contribution of plant–consumer interactions within local food webs and weakening herbivore-mediated energy pathways near roads.

Our structural equation models identified an indirect pathway linking road proximity to reduced herbivory: sites closer to roads supported lower herbivore richness, which in turn reduced total herbivore biomass and ultimately weakened modelled herbivory flux. This pattern may reflect the broader role of roads as proxies for landscape accessibility and cumulative anthropogenic pressure, rather than the local effects of roads alone. Increased accessibility can facilitate hunting ^39^, habitat degradation ^6^, deforestation ^15^, and changes in species interactions ^38^, all of which may disproportionately affect herbivorous vertebrates, including large-bodied game species such as pacas, agoutis, peccaries, brocket deer, tinamous, tapirs, and curassows ^13,40^. Although mean body size did not directly vary with road distance in this study, reductions in herbivore richness and associated declines in biomass were sufficient to weaken herbivory fluxes. This pattern reflects a contraction of herbivore communities rather than any shift in their size structure: fewer species and fewer individuals simply assimilate less energy from plant resources, reducing the overall amount of plant energy processed. Thus, the indirect pathway—operating through reductions in herbivore richness and biomass—represents a major mechanism by which road proximity depresses plant-to-vertebrate energy flows, independently of changes in size distributions.

Together, these findings suggest that roads may reduce herbivory through both indirect effects on community structure and broader functional reorganization of trophic interactions. Herbivory and carnivory are tightly interconnected functions, and the opposing responses we observed may indicate the emergence of trophic cascades along disturbance gradients that need to be better investigated. Higher levels of carnivory near roads coincided with lower herbivory, suggesting a functional reorganisation of food webs along the disturbance gradient. Previous work in tropical systems has shown that land-use change can reduce trophic interactions and diminish energy transfer to higher trophic levels ^41^. In contrast, our results suggest an opposite configuration under low-impact Amazonian conditions: more energy is channeled into predators and less into herbivores in road-adjacent environments. This pattern may represent an early-stage trophic imbalance, where generalist predators can benefit from the resource conditions near roads and altered resource conditions before stronger biodiversity losses become evident. Our findings suggest that under higher levels of human pressure, such as those found along the arc of deforestation, the shifts on ecosystem functions and conventional biodiversity metrics (e.g. species richness and abundance) may become even more pronounced.

Finally, our findings highlight the importance of considering trophic interactions into assessments of ecosystem functioning. Vertebrate functional groups, including frugivores, browsers, granivorous ungulates, primates and predators, play crucial roles in the forest regeneration dynamics by dispersing seeds, shaping vegetation structure, and regulating herbivore populations, preventing overbrowsing and maintaining trophic balance ^42^. Together, these processes sustain forest recovery and ecosystem resilience after disturbance. While some species can continue to persist in landscapes with limited human impacts, our study focuses on the broader, landscape-scale consequences of expanding road networks and associated accessibility, which can amplify pressures on vertebrate communities and disrupt the energy pathways that sustain ecosystem functioning. By showing that subtle shifts in energy flows emerge even under relatively mild degradation, our energetic approach reveals early warning signals of ecosystem disruption that precede detectable losses in species richness or abundance. As habitat alteration and physical accessibility expand across the Amazon, through new road networks and advancing agricultural frontiers, safeguarding both species and the energetic pathways connecting them will be essential to preserve ecosystem functioning and ensure the resilience of Amazonian forests in the decades to come.

Our study highlights the importance of assessing ecosystem functioning through multiple metrics, with the energetic approach offering a powerful tool to quantify functions provided by animal communities ^29,41,43^. Although many resilient taxa—including key seed dispersers such as large frugivorous birds, ungulates, and primates—can persist under intermediate disturbance and help sustain forest recovery when connectivity is maintained, their presence does not fully buffer the broader reconfiguration of trophic processes associated with road proximity. Nevertheless, these species remain crucial for post-disturbance regeneration: many tolerate low-intensity habitat modification and continue to support recovery processes, likely facilitated by low hunting pressure and the connection of forest remnants to intact forest. These changes may also have broader implications for forest resilience through cascading effects on plant–animal interactions, particularly as recent evidence suggests that disturbed Amazonian forests may undergo processes of ecological homogenization and increasing dominance of generalist tree species ^44^. As infrastructure continues to expand across the Amazon, safeguarding both vertebrate communities and their energy pathways in roadless areas will be essential to preserve ecosystem functioning ^45^ and sustain the ecological resilience of these forests.

## 2. METHODS

### Study area

Our study covers a network of camera-trap clusters located throughout Amazon forests as defined by the RAISG (Amazonian Network of Georeferenced Socioenvironmental Information) phytogeographic and hydrological boundaries, representing the world’s largest continuous tropical forest. The region is structured by large river systems, pronounced rainfall gradients of 1,400–4,500 mm annually ^46^, and contrasting geomorphological zones ranging from upland unflooded (terra firme) forests to extensive wetlands. Floodplain areas differ markedly in hydrology and sediment load, resulting in distinct forest types shaped by fluvial geochemistry, including white-, clear-, and black-water rivers ^47^. These environmental gradients create a heterogeneous macrohabitat mosaic that supports exceptionally rich vertebrate communities. Accordingly, the camera-trap sampling sites included in this study span a wide spectrum of environmental gradients and encompass both protected and unprotected areas. A full description of the study region, ecosystem types, and sampling sites is available in ^25^.

### Dataset standardization and species trait harmonization

We compiled species records from the Amazonia Camtrap dataset ^25^, which includes vertebrate species sampled using camera traps across multiple Amazonian study areas. We retained only records of wild mammals and birds, excluding rare records of domesticated species. Because our focus is on trophic interactions, we excluded datasets that targeted a single taxonomic group. We also excluded datasets from cameras installed in the forest canopy due to their limited representation among all available studies. We retained only medium- to large-bodied vertebrates (≥400g for birds, and ≥900g for mammals), which are more consistently detected by camera traps. Taxonomic details were resolved to the species level and standardized using GBIF’s nomenclature backbone for consistency.

To reconstruct the trophic network and evaluate the fluxes among species within the Amazonia, we extracted diet composition and species body mass from the EltonTraits database ^48^, which were used to infer trophic links and estimate metabolic demand. As these values represent standardized species-level traits, local intraspecific variation across populations was not explicitly accounted for. Based on the diet data, we classified each species into functional groups to quantify community attributes for both carnivores and herbivores (Supplementary Fig 2). Species were assigned to a broad carnivore trophic guild if they consumed any proportion of animal prey, including invertebrates and vertebrates, or to a herbivore guild if they consumed any proportion of plant material, including leaves, fruits, seeds and nectar ^26,27^. These guild definitions allowed species to contribute to multiple trophic pathways in omnivores, whose diet included both plant and animal material.

Using data from the EltonTraits database, we removed five vulture species classified as strict scavengers (≥70% necrophagous diet) and retained only bird species that forage below the canopy (i.e., terrestrial or near-ground foragers), defined as species with ≥50% of their foraging activity on the ground or up to 2m height. This restriction was applied to reduce detection bias, as our ground-based camera trapping design is less effective in sampling arboreal species. Missing trait values were filled using the Avonet database (for birds), Pantheria database (for mammals), and genus-level trait means (for which species-level data were unavailable).

### Definition of sampling units (camera-trap clusters)

We defined 85 sampling units by spatially clustering individual camera-trap stations into groups representing local food webs. Clusters were defined based on three criteria: (i) whenever possible, cameras from the same data provider were retained within the same unit; (ii) the maximum distance between any two cameras in a cluster did not exceed 15 km; and (iii) stations located on opposite sides of major rivers were grouped separately, as these rivers represent major ecological and physical barriers for most terrestrial mammals and birds. Major rivers were identified through visual inspection of regional hydrographic layers in QGIS (version 3.6,0; QGIS Development Team) and correspond to the main river channels across the study region. Cluster boundaries were delineated using the minimum convex hull of its constituent camera-trap stations. All clustering decisions were implemented manually in QGIS, integrating spatial proximity with landscape context.

### Distance to Roads

Road data were obtained from the RAISG road dataset ^49^, which compiles the best available road data across Amazonian countries and provides one of the most consistent harmonized basin-wide road datasets currently available for the region, including paved highways as well as secondary and unpaved roads. Given that both paved and unpaved roads are known to drive ecological degradation ^50^, we grouped them together in our analysis. Planned roads that had not yet been built were excluded. However, because informal or temporary roads may be underrepresented, distance-to-road estimates may contain some spatial measurement uncertainty. We calculated the Euclidean distance from the centroid of each cluster to the nearest road. Because clusters represent the unit of analysis, centroid distance provided a standardized measure of road proximity at the scale of the sampled community. Final distances were expressed in kilometers and used as a cluster-level covariate.

### Construction of the food webs

To quantify trophic functions associated with energy channels (carnivory and herbivory), we reconstructed local food webs among all the species recorded by camera traps within each cluster across the Amazon. To predict potential trophic interactions, we first parameterized the size-constrained feeding niche (SCFN) model ^51^ using a compiled empirical food-web dataset that included mammal and bird species from ^52,53^. We manually reviewed and validated all predicted links (2,234) using available dietary references and long-term expert knowledge from C.A.P. on Amazonian vertebrate ecology, resulting in a curated metaweb of potential interactions for Amazonian mammals and birds (provided as Supplementary data 1). Details on food-web construction are provided in Supplementary Information (Section 2). From this metaweb, we generated cluster-specific interaction networks (local food webs) for each camera-trap cluster, represented as binary prey–predator matrices which were complemented with three basal resource nodes (plants, carrion, and live animal prey) to ensure trophic completeness.

### Relative biomass estimation

To estimate the energy required to sustain trophic networks, we calculated species biomass at each cluster by multiplying relative abundances by the average body mass. To estimate species relative abundances across camera-trap clusters, we used a hierarchical multi-species N-mixture model that jointly accounts for imperfect detection from the camera records and variation across sites ^54–57^. For each species, true abundance at each site was modelled as a (zero-inflated) Poisson process influenced by environmental covariates (road distance), while observed counts arose from a Binomial detection process with survey-level covariates affecting detection probability (camera trapping effort, time between detections, baited or unbaited). Species-specific responses to both abundance and detection covariates were treated as random effects, ensuring partial pooling of information across species and improving estimation accuracy, especially for rare taxa. A detailed explanation of the model and its assumptions can be found in the Supplementary Information (Section 3).

To ensure comparability across species, relative abundances were rescaled using an allometric correction based on body mass, thereby accounting for increased detection probability associated with the larger movement ranges of larger species. Specifically, abundances were divided by M^0.75^, reflecting the scaling between body mass and ecological rates linked to metabolism and movement ^58,59^. This adjustment standardizes abundances to a common area, yielding relative, area-corrected abundance estimates that can be compared across species and used in biomass calculations. Species-level biomass was estimated by multiplying relative abundance by body mass. To estimate trophic biomass at the cluster level, we summed species-level biomasses across all species contributing to each trophic niche. Species contributing to carnivory were defined as those with non-zero animal consumption, whereas species contributing to herbivory were defined as those with non-zero plant consumption.

### Flux evaluation

Energy fluxes were quantified by combining A) cluster-specific trophic interaction networks, B) body mass–based metabolic rates, C) prey-specific assimilation efficiencies, and D) species biomass estimates, as summarized in Fig. 5. For this we used the *fluxweb* package in R ^60^. We calculated the total fluxes to herbivores (fluxes from plants to vertebrates) and carnivores (fluxes from prey to predators) per cluster (local food webs) to assess herbivory and carnivory functions, respectively. To assess the energy required to sustain functioning of the species, we estimated per-biomass metabolic rates using order-specific allometric equations based on species body mass, with coefficients derived from empirical data for endothermic vertebrates reported by ^61^. We defined assimilation efficiencies as the proportion of consumed energy assimilated by the consumer, assigning values at the prey level according to dietary type, following empirical values from ^62,63^: 70% from plants, 80% from animal prey, and 85% from carrion. For resources not detected by camera traps, we added three basal resource nodes to every food web: plants, animal prey (invertebrates and small vertebrates not recorded by cameras), and carrion. Diet information from EltonTraits ^48^ was used to assign each consumer the proportion of its diet derived from these resource categories. After assembling the interaction matrix and matching it to species-specific biomass and metabolic rates for each cluster, we allocated each consumer’s total energy intake across its potential food sources. This allocation relied on two complementary sources of information. First, for animal prey detected by camera traps, feeding preferences were computed using the biomass-weighted allocation implemented in *fluxweb*, which distributes a predator’s consumption among prey in proportion to their relative biomass. Second, for resources not captured by camera traps—such as plants, invertebrates, and small vertebrates—we used the consumer’s diet composition to allocate the remaining fraction of its intake among these food categories. Together, these steps ensured that omnivores partitioned their consumption rates across all potential food sources in a way that reflects both their intrinsic dietary preferences and prey availability in the local community.

**Fig. 5:**
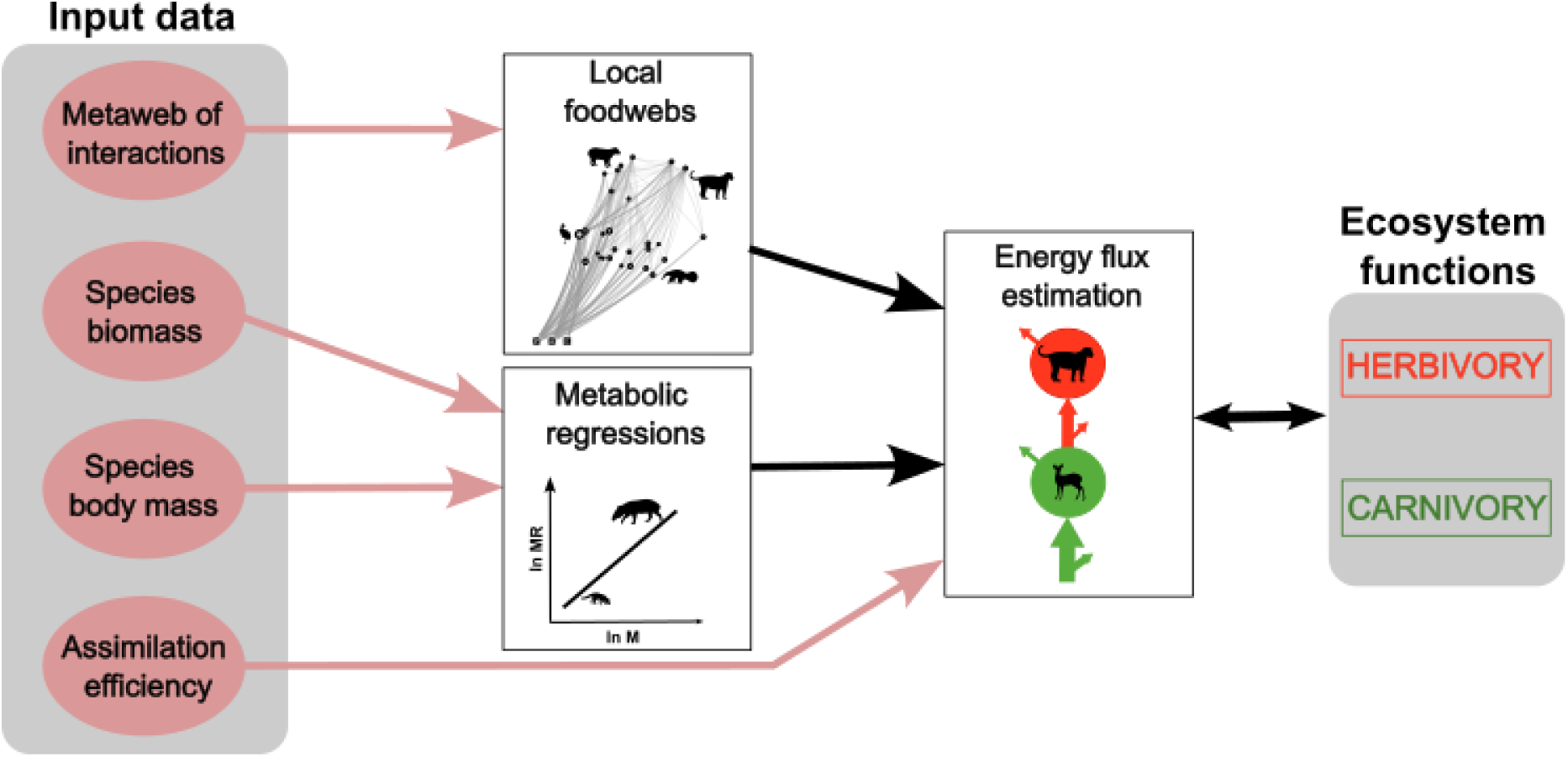
Conceptual framework for estimating ecosystem functions. Local food webs are derived from a metaweb of potential trophic interactions. Species body mass is used to estimate metabolic rates via allometric relationships, which are scaled by species biomass to quantify the energetic demand of each node. Network structure, metabolic losses, and assimilation efficiencies are integrated within the fluxweb framework to calculate energy fluxes across trophic interactions, from higher to lower trophic levels. These fluxes are then aggregated to quantify ecosystem functions, specifically herbivory (plant-to-consumer flux) and carnivory (consumer-to-consumer flux).

To characterize energy flow within each local food web, we calculated the realized flux matrix assuming a steady-state energy balance. To characterize ecosystem trophic functions, we extracted herbivory and carnivory from the flux matrices by summing the outgoing fluxes from plants and animals. Herbivory captured all plant-derived energy assimilated by vertebrate herbivores, while carnivory captured all animal prey-derived energy assimilated by vertebrate carnivores (excluding carrion, which represents a distinct function). Because both abundances and energy fluxes are expressed relative to an effective sampling area (as implemented in *fluxweb*), our final flux estimates are unitless and should be interpreted as relative rates of energy flow rather than absolute energetic quantities. These values quantify relative per-area consumer energetics, providing unitless measures of trophic function.

### Data analysis

We built the path diagram including hypothesized direct effects of community characteristics (species richness, biomass and mean body mass), and indirect effects of distance to roads through the community characteristics on ecosystem functions (carnivory and herbivory) (Supplementary Figure 3). To test the effects of distance to nearest roads on energy fluxes, prior to analyses, all continuous and positive variables were log₁₀-transformed to improve variance homogeneity, including distance to road, total biomass, mean body mass, species richness, and trophic fluxes. We then applied a piecewise structural equation modelling (SEM) framework based on confirmatory path analysis ^64^ to evaluate how road distance influences carnivory and herbivory fluxes, both directly and indirectly through changes in species richness, mean body mass, and biomass. Separate SEMs were fitted for the ecosystem functions of carnivory and herbivory. Herbivory and carnivory represent distinct trophic pathways, involving partly overlapping species subsets and potentially opposing responses to road proximity. Modelling them in separate but structurally analogous SEMs allowed us to address our main question—how roads affect each function—while avoiding unnecessary complexity and untested assumptions about direct causal links between herbivory and carnivory at the guild level. In the initial model, road distance was specified as a direct driver of species richness and mean body mass, and these two variables—together with road distance—were modelled as predictors of community biomass. Energy flux was modelled as a direct function of biomass, mean body mass, and species richness, under the hypothesis that road effects on trophic fluxes are mediated through shifts in these community attributes rather than acting directly on fluxes. Model alternatives were compared using Fisher’s C and directed separation tests, which indicated that a direct path from road distance to flux was missing, whereas the path from road distance to biomass was not supported. Final SEMs therefore retained the direct road–flux relationship while excluding the road–biomass path. To account for spatial autocorrelation among clusters distributed across the Amazon basin, Gaussian submodels (fluxes, biomass, and mean body mass) were fitted using generalized least squares (GLS) models with an exponential spatial correlation structure based on latitude and longitude. Final SEMs thus integrated spatial GLS and spatial Poisson GLMM submodels within a unified piecewise SEM framework.

## ACKNOWLEDGEMENTS

A.C.A. acknowledges support from the working group Theory in Biodiversity Science, and by sDiv, the Synthesis Centre for Biodiversity Sciences of iDiv, funded by the German Research Foundation (DFG – FZT 118, 202548816). A.B.C. acknowledges a doctoral scholarship from the Conselho Nacional de Desenvolvimento Científico e Tecnológico (CNPq). H.M.P. and J.H. acknowledge funding from iDiv through the German Research Foundation (DFG – FZT 118, 202548816). R.B.W. acknowledges support from Wildlife Conservation Society country programs across the Amazon basin. R.V.M. and R.d.P.R.G. acknowledge support from Conservación Internacional – Team Network.

## AUTHOS CONTRIBUTIONS

A.C.A., U.B., B.G. and E.B. conceived and designed the study. A.C.A., U.B., B.R., B.G., E.B., J.L., C.A.P., H.M.P., C.M., J.H., A.M. and T.S. contributed to the conceptual development of the study and participated in discussions throughout the project. B.R. developed the abundance models. J.L. developed the predator–prey interaction model. C.A.P. reviewed and validated trophic interactions among species. A.C.A. assembled the datasets, conducted the analyses, and wrote the first draft of the manuscript. All remaining authors contributed to data collection, curation, and/or provision of biodiversity records. All authors reviewed, edited, and approved the final manuscript.

