## Supplementary Information for "Forest degradation reshapes trophic functioning in vertebrate food webs across Amazonian forests"

SUPPLEMENTARY FIGURES

**A**

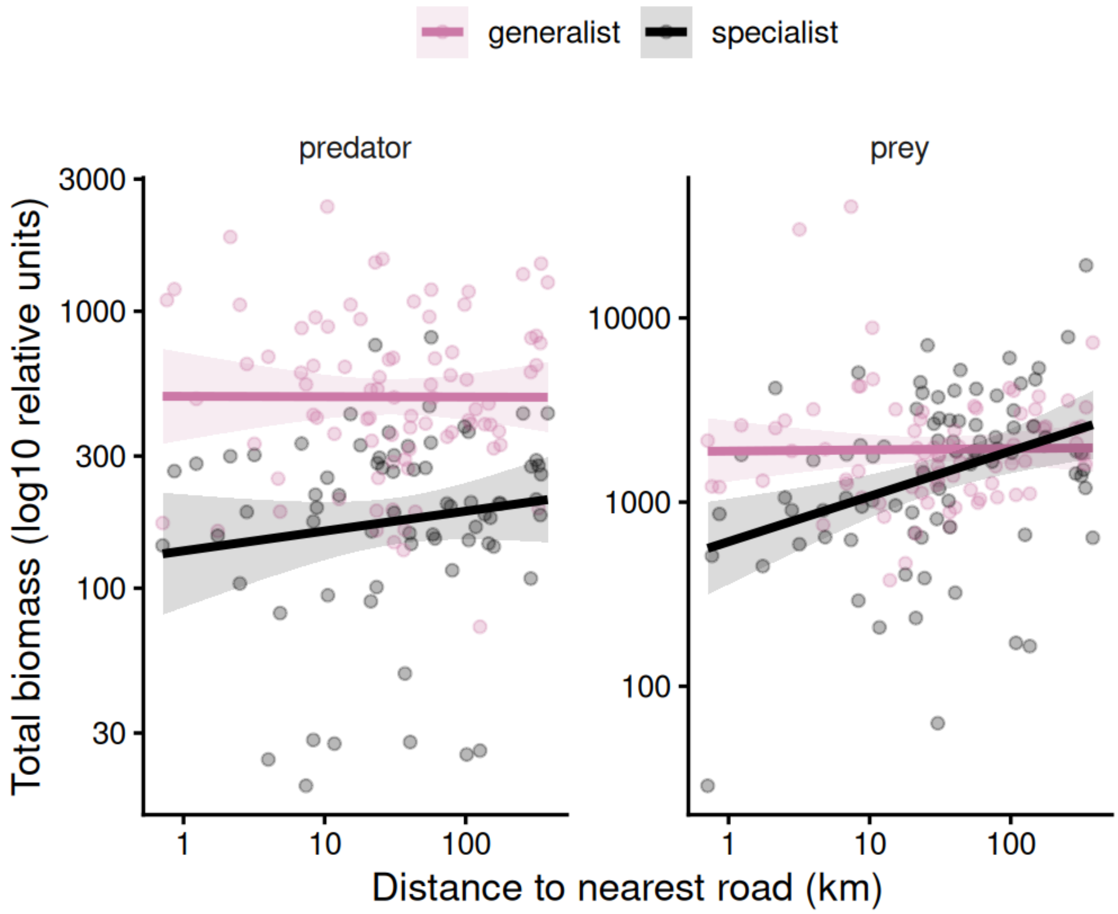

**B**

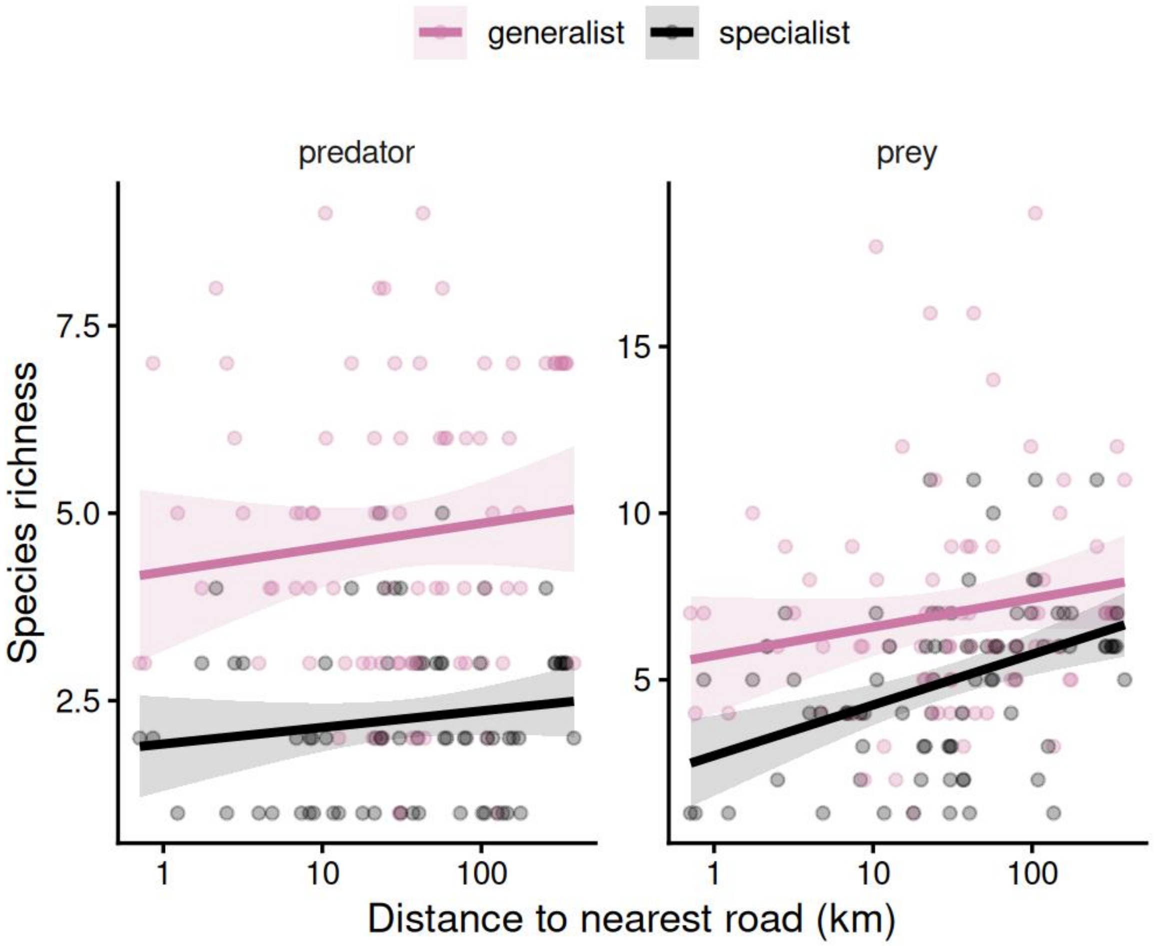


***Supplementary Fig 1. Biomass and species richness of generalist and specialist species along a road-distance gradient.***

*Relationships between distance to the nearest road and (A) total biomass and (B) species richness for predators and prey. Points represent individual clusters, and solid lines indicate 95% confidence intervals. Colors distinguish generalist (pink) and specialist (black) species. Species were classified as generalists or specialists following Prist et al. 2012 when available, and complemented using diet information from EltonTraits, where species with ≥80% reliance on a single diet category were classified as specialists, and all others as generalists.*

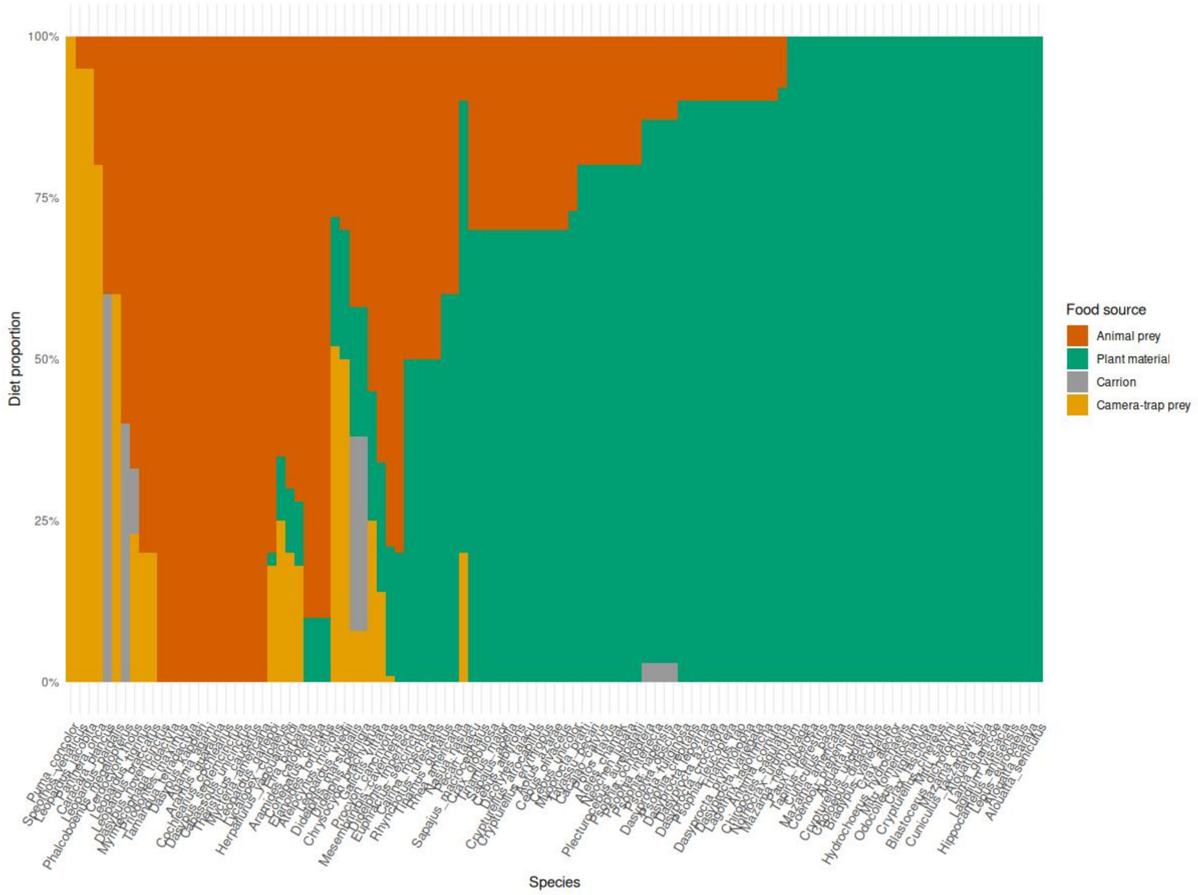


***Supplementary Fig 2. Diet composition across vertebrate species included in the analysis.*** *Each bar represents a species and shows the proportional contribution of different food sources to its diet: plant material (green), animal prey (orange), camera-trap-detectable vertebrate prey from Antunes et al. 2022 (yellow), and carrion (grey).*

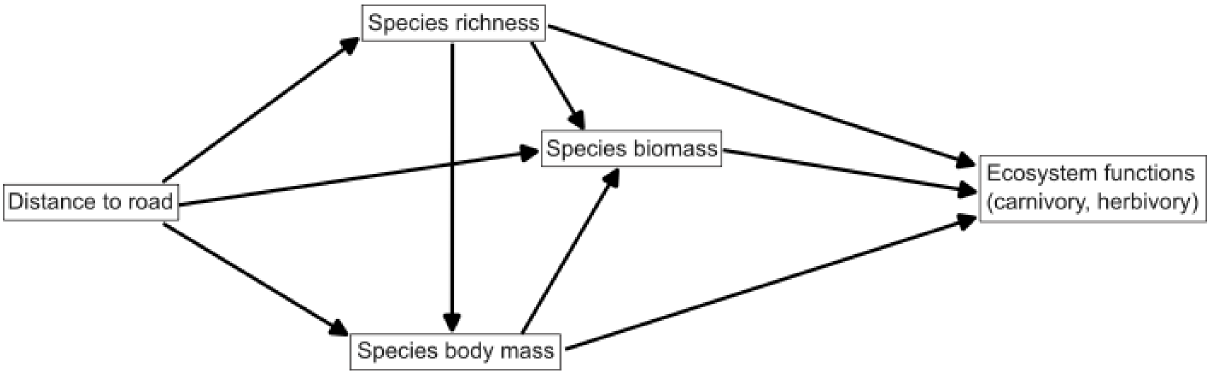


***Supplementary Fig 3.*** **A-priori structural equation model.** *Path diagram of hypothesized direct and indirect effects of distance to roads on ecosystem functions (carnivory and herbivory fluxes).*

SUPPLEMENTARY INFORMATION (SECTION 1)

One cluster included in the analysis is located in Yasuní National Park, an Ecuadorian region impacted by road development linked to oil extraction and home to Waorani Indigenous communities. We considered this site as an outlier because, although it lies in close proximity to a road, this road primarily serves oil operations and does not connect to broader road networks. As a result, the site can still be considered as remote and spatially isolated compared to the other clusters included in our analysis. Despite relatively low traffic, this context should be interpreted with caution, as low traffic does not necessarily imply low ecological impact. Previous local studies have documented a depletion of the local wildlife populations (including jaguars, as well as game species such as tinamous and peccaries), and a two-fold increase of the bushmeat extraction area, potentially threatening the livelihoods of local indigenous groups (Espinosa et al. 2018; Espinosa et al. 2014; Suárez et al. 2013). Here, our objective is not to evaluate the impact of individual roads, but rather to capture broad-scale patterns in ecosystem functioning across Amazonia. To evaluate the influence of this potentially distinct site, we repeated all analyses with and without this cluster and obtained qualitatively consistent results.

Altogether, we sampled 86 camera-trap clusters, detecting 35 bird and 72 mammal species. Of these, 79 species contributed to carnivory fluxes and 85 to herbivory fluxes, with 57 species contributing to both trophic pathways. For carnivory, the final SEM (Fig. 3a) showed an adequate fit (Fisher's C = 0.70, $p = 0.70$). Carnivory fluxes increased with carnivore biomass (standardized coefficient $\beta = 0.27$, $p = 0.015$) and carnivore richness ($\beta = 0.54$, $p < 0.001$), while the effect of mean body mass on fluxes was small and non-significant ($\beta = 0.05$, $p = 0.48$). Importantly, road distance had a negative direct effect on carnivory flux ($\beta = -0.19$, $p = 0.017$), indicating higher levels of carnivory nearer roads after accounting for richness, biomass, and body mass. Carnivore biomass was strongly and positively related to richness ($\beta = 0.81$, $p < 0.001$) and negatively related to mean body mass ($\beta = -0.17$, p =

0.014), whereas road distance had no detectable effect on carnivore richness (β = 0.14, p = 0.40) or mean body mass (β = −0.09, p = 0.48).

For herbivory, the final SEM including all 86 clusters (Fig. 3b) also showed a good fit (Fisher's C = 0.85, p = 0.85). Herbivory fluxes increased strongly with herbivore biomass (β = 0.53, p < 0.001) and, to a lesser extent, herbivore richness (β = 0.23, p = 0.043), whereas the effect of mean body mass was small and non-significant (β = −0.001, p = 0.99). Road distance had a positive direct effect on herbivory flux (β = 0.20, p = 0.017), indicating higher herbivory farther from roads. Herbivore biomass increased strongly with richness (β = 0.78, p < 0.001), while road distance showed no detectable effect on herbivore richness (β = 0.22, p = 0.25) or mean body mass (β = 0.07, p = 0.53). Overall, these results suggest that herbivory patterns were primarily structured by internal community properties, particularly richness and biomass, with an additional positive direct association with distance to roads.

SUPPLEMENTARY INFORMATION (SECTION 2)

*Construction of food webs in Amazon with a size-constrained feeding-niche (SCFN) model*

To predict the potential trophic interaction between the species recorded by the camera traps in the Amazon, we first parameterized the size-constrained feeding-niche (SCFN) model (eq.I-IV) by fitting it with a compiled food-web dataset (mammal and bird species from TETRA-EU 1.0 and Caron et al. 2024) using the Bayesian method. We excluded duplicate interaction records to eliminate density effects. The compiled food-web dataset included both feeding and non-feeding interactions information between species. Specifically, for a given species x, interactions with its prey j and predators i were coded as 1 (i.e., L_ix = 1, L_xj = 1), whereas the non-feeding interactions were coded as 0 (i.e., L_ix = 0, L_xj = 0). This completion of the dataset was necessary for fitting SCFN model (eq. I) using the Bayesian method as follows:

$$p_{ij} = \theta_i \, exp\,(-\frac{(m_i - m_j - \mu_i)^2}{2{\sigma_i}^2}) \quad \text{(eq.I)}$$

where $p_{ij}$ is the feeding probability of predator i feeding on prey j, $m_x$ is the log_10 body sizes of species x, $\mu_i$ is the log_10 optimal size ratio of predator i, and ${\sigma_i}^2$ represents the feeding range of predator i. $\theta_i$ is the maximum feeding probability of predator i. The three characteristics of the size-constrained feeding niche can be expressed as functions of predators body mass $m_i$:

$$\mu_i = \beta_{\mu,0} + \beta_{\mu,1} m_i \quad \text{(eq.II)}$$

$${\sigma_i}^2 = exp\,(\beta_{\sigma^2,0} + \beta_{\sigma^2,1} m_i) \quad \text{(eq.III)}$$

$$\theta_i = \frac{exp(\beta_{\theta,0} + \beta_{\theta,1} m_i)}{1 + exp(\beta_{\theta,0} + \beta_{\theta,1} m_i)} \quad \text{(eq.IV)}$$

The compiled food web dataset is composed of 4 food webs with 1,029 species (mammals and birds) and 28,931 realized interactions records. For each food web, we extracted all prey and predator species and constructed an adjacency matrix to characterize the trophic network. We fitted the SCFN model using the Bayesian method with Stan (version 2.26.1) in R (version 4.2.3). We used non-informative priors for all 6 parameters, and we ran the Markov chain Monte Carlo sampling for 16,000 iterations with 8,000 warm-ups. The posterior distributions for the six parameters that describe the three characteristics of the feeding niche have been recorded in Table 1. We fitted the SCFN model for birds and mammals separately. Based on the estimated feeding-niche parameters (that is, $\mu_i$, $\sigma_i$, and $\theta_i$), we calculated the feeding probability $p_{ij}$ for all possible predator–prey relationships among Amazonian vertebrates using information on species body mass and dietary data.

| **parameter** | **posterior distribution** |
|---|---|
| $\beta_{\mu,0}$ | 7.467 (7.067, 7.878) |
| $\beta_{\mu,1}$ | -0.486 (-0.529, -0.443) |
| $\beta_{\sigma,0}$ | 1.625 (1.533, 1.717) |
| $\beta_{\sigma,1}$ | 0.126 (0.116, 0.137) |
| $\beta_{\theta,0}$ | -2.579 (-2.703, -2.454) |
| $\beta_{\theta,1}$ | -0.146 (0.130, 0.163) |

***Table 1. posterior distribution of the parameters in the SCFN model.***

To convert the probability $p_{ij}$ into presence–absence information (Lij=1L_{ij} = 1Lij=1 or 000), we considered that species i is a predator of species j if $p_{ij}$ is larger than a pre-assigned threshold $p^*$. The threshold $p^*$ was defined as a randomly assigned value between zero and the maximum $p_{ij}$ observed in this study. Species identified in the literature as herbivores were restricted to being prey only, meaning they could only be eaten by other species, and cannibalistic interactions were excluded.

Because the size-constrained feeding niche may not be applicable to all species, we manually reviewed and validated all predicted interactions after the simulation. Specifically, we checked all 2,234 predicted interactions using available dietary references and expert knowledge from C.P. on Amazonian vertebrate ecology. If a predicted interaction ($L_{ij} = 1$) was not supported by available knowledge or ecological plausibility (for example, incompatible diets or unrealistic predator–prey size relationships), the interaction was removed ($L_{ij} = 0$). Conversely, if an interaction predicted as absent ($L_{ij} = 0$) was documented in the literature, it was added to the network ($L_{ij} = 1$). This procedure resulted in a curated metaweb of potential trophic interactions among Amazonian mammals and birds (Supplementary Data 1).

SUPPLEMENTARY INFORMATION (SECTION 3)

*Abundance estimation*

We constructed a multispecies N-mixture model (Doser et al. 2024, Kery & Royle 2016) in R-Nimble to estimate abundances $N_{ij}$ of species *i* in location *j* based on observed counts $y_{ijk}$ from a location *j*'s cameras *k*. The **abundance equations** are

$$N_{ij} \sim Poisson(\mu_{ij}\psi_{ij})$$

$$\psi_{ij} \sim Bernoulli(\phi_i)$$

$$log(\mu_{ij}) = \beta_{0i} + \beta_{1i}x_{1j}$$

where $\psi_{ij}$ is a discrete latent variable (0 or 1) for occupancy and $\phi_i \in (0,1)$ is a species' mean occupancy across locations, defining a zero-inflated Poisson model (Ponisio et al. 2020). $\mu_{ij}$ is the GLMM prediction for the single, location-dependent predictor $x_{1j}$ with species-dependent intercepts $\beta_{0i}$ and slopes $\beta_{1i}$. The estimated abundances $N_{ij}$ are connected to camera counts $y_{ijk}$ via the **detection equations**

$$y_{ijk} \sim Binomial(p_{ijk}, N_{ij})$$

$$logit(p_{ijk}) = \alpha_{0i} + \alpha_1 x_{1jk} + \alpha_2 x_{2jk} + \alpha_{3i} x_{1jk}$$

where $p_{ijk}$ is the GLMM prediction for detection probability of an individual for a specific camera trap. Predictors are camera effort $x_{1jk}$, a camera's time between two separate detections $x_{2jk}$ and bait $x_{3jk}$. The first two predictors have species-independent effects $\alpha_1$ and $\alpha_2$, while intercepts $\alpha_{0i}$ and effects of bait $\alpha_{3i}$ are species-specific. For both GLMMs, a **partial pooling** approach was used with species-specific effects assumed to be normally distributed

$$\beta_{0i} \sim Normal(\beta_{0\mu}, \beta_{0\sigma})$$

$$\beta_{1i} \sim Normal(\beta_{1\mu}, \beta_{1\sigma})$$

$$\alpha_{0i} \sim Normal(\alpha_{0\mu}, \alpha_{0\sigma})$$

$$\alpha_{3i} \sim Normal(\alpha_{3\mu}, \alpha_{3\sigma})$$

with joint means and standard deviations $\beta_{0\mu}$, $\beta_{0\sigma}$ etc. We used weakly regularizing standard-normal **priors** for all unrestricted model parameters ($\beta_{0\mu}$, $\beta_{1\mu}$, $\alpha_{0\mu}$, $\alpha_1$, $\alpha_2$, $\alpha_{3\mu}$), half-normal priors for positive parameters ($\beta_{0\sigma}, \beta_{1\sigma}, \alpha_{0\sigma}, \alpha_{3\sigma}$), and uniform (0,1) priors for all $\phi_i$.

For MCMC model fitting we used 3 chains with $10^6$ iterations each, the first half of the iterations used for the burn-in phase.
